# Hierarchical cue control of drug seeking in the face of cost

**DOI:** 10.1101/2022.05.05.490799

**Authors:** Val Collins, Kaisa N. Bornhoft, Amy Wolff, Sonal Sinha, Benjamin T. Saunders

**Affiliations:** Department of Neuroscience, University of Minnesota; Medical Discovery Team on Addiction, University of Minnesota; University of California, San Francisco; Johns Hopkins University

## Abstract

Addiction is characterized by intermittent drug seeking despite rising costs. This behavior is heavily influenced by environmental stimuli that signal drug availability and reinforce seeking. We aimed to establish the relationship between three key aspects of human drug use in rats: the intermittent, binge nature of drug intake, the motivational conflict of drug seeking in the face of escalating negative costs, and the ability of different drug cues to interact to modulate relapse. Rats were trained to self administer cocaine on an intermittent access schedule, where brief drug availability states were signaled by a shift in the ambient lighting of the environment and cocaine delivery was signaled by a separate proximal cue. Rats then went through a conflict procedure, where foot shock intensity associated with cocaine seeking was escalated until intake was suppressed. We completed relapse tests where the drug delivery cue was non contingently presented alone, or in the context of dynamic drug availability state transitions. Intermittent access spurred psychomotor sensitization and binge-like cocaine intake. The intensity of binge-like drug taking during training was predictive of later drug seeking despite escalating costs. In relapse tests, the ability of a proximal drug cue to trigger relapse was gated by the presence of a global cue signaling drug-availability state transitions. Our results suggest that the pattern of drug intake plays a role in many features of addiction, including modifying an individual’s willingness to endure high costs associated with drug seeking. Further, our results indicate that drug-related sensory information can be hierarchically organized to exert a dynamic modulating influence on drug-seeking motivation.

## Introduction

Addiction is characterized by intermittent drug seeking despite rising costs, with a persistent threat of relapse (Robinson and Berridge 1993; Shaham et al. 2003; Everitt and Robbins 2005; Poisson et al. 2021). This behavior is heavily influenced by environmental stimuli that signal drug availability and receipt of drugs to invigorate and reinforce drug seeking actions. Drug-related cues take on many forms, including focal, proximal cues and broader global cues signaling drug-related contexts and states (Chaudhri et al. 2008; Crombag et al. 2008; Fraser and Holland 2019).

A growing body of preclinical work has made use of intermittent access schedules that promote rapid spikes in brain drug levels, which uniquely drive exaggerated motivation and striatal plasticity that is important in addiction (Zimmer et al. 2012; Bentzley et al. 2013; Calipari et al. 2013; Kawa et al. 2019b; Allain and Samaha 2019; Carr et al. 2020; Samaha et al. 2021). Earlier studies demonstrated that the pharmacokinetics of drugs, rather than the absolute amount of a drug, powerfully determines their ability to engage brain reward systems and behavior (Budney et al. 1993; Cone 1995; Samaha et al. 2002, 2005; Allain et al. 2015; Kawa et al. 2019a; Canchy et al. 2021). Drug cues are centrally positioned to modulate drug seeking motivation and, in turn, the pattern of drug intake. Notably, there are considerable individual differences in the extent to which drug-related sensory information motivates behavior (Flagel et al. 2009; Robinson et al. 2014). In human substance use disorder (SUD) patients, the interplay of drug access patterns, dynamic environmental stimuli, drug seeking costs, and individual vulnerabilities produces a difficult landscape for effective treatments (Bertz et al. 2018; Venniro et al. 2020).

Here, we explored the intersection of drug cues, costs, and individual differences in rats. We integrated behavioral tasks that capture three key features of addiction: 1) the volitional, intermittent, binge-and-stop nature of drug use, 2) continued use despite escalating costs, and 3) the interaction of different levels of drug-related sensory information in the control of relapse. We used an intermittent-access cocaine self-administration paradigm that leads to escalating, rapid drug intake that more closely models human drug use compared to traditional continuous access models (Allain et al. 2015; Kawa et al. 2019a). Additionally, we used a conflict paradigm to assess motivation to continue drug taking in the face of escalating foot shock (Cooper et al. 2007; Saunders et al. 2013). Lastly, we compared the effect of noncontingent presentations of global drug availability cues and proximal drug delivery cues to promote relapse.

Our results motivate three main conclusions. First, sensory information signaling drug availability can dynamically regulate drug seeking in part via hierarchical modulation of the motivational value of other drug-related cues. Second, rapid, intermittent cocaine intake patterns produce cost insensitivity in future drug seeking. Third, there are considerable individual differences in the extent to which rats use state-level/global versus discrete/proximal drug cues to guide drug seeking, suggesting unique trajectories to relapse. Together, our results underscore that the complex set of variables, including individual decision-making strategies, hierarchically organized drug-associated stimuli, dynamic patterns of access to drugs, and the cost of drug seeking, interact to regulate drug seeking motivation and addiction-like behaviors.

## Results

### Intermittent access produced binge-like drug taking invigorated by drug-availability state onset

Rats first learned to self-administer cocaine in a continuous access paradigm with a FR1 schedule of reinforcement (Supplemental Figure 1B). By the final day of acquisition, animals learned to discriminate between the active and inactive nose ports (Supplemental Figure 1C; Multiple paired t-test, t(32)=p<0.0001, n=35). Males and females used similar amounts of cocaine throughout acquisition (Supplemental Figure 3A; 2-way ANOVA, main effect of sex F(5,97) = 1.929, p=0.0964). Following the 3-day acquisition of continuous access cocaine self-administration, rats underwent 14 days of intermittent access. Within this task (Fig.1B), brief 5-min periods of cocaine access were signaled by a drug-availability state cue (turning off all ambient lights illuminating the chamber). Interspersed were longer periods where drug was unavailable - these no-drug periods were signaled by illumination of the chamber. During the drug-availability periods, rats could nose poke for a 3.2-sec cocaine infusion via an in-dwelling i.v. catheter that coincided with the presentation of a 3.2-sec drug-delivery cue (illuminated nose port and white noise). Behavior on days 1 and 14 is shown in Figure 1C. Males and females showed similar patterns (Supplemental Figure 3B). Responding in the unpaired group was overall low and showed no clear pattern (Supplemental Fig. 2B).

**Figure 1.**
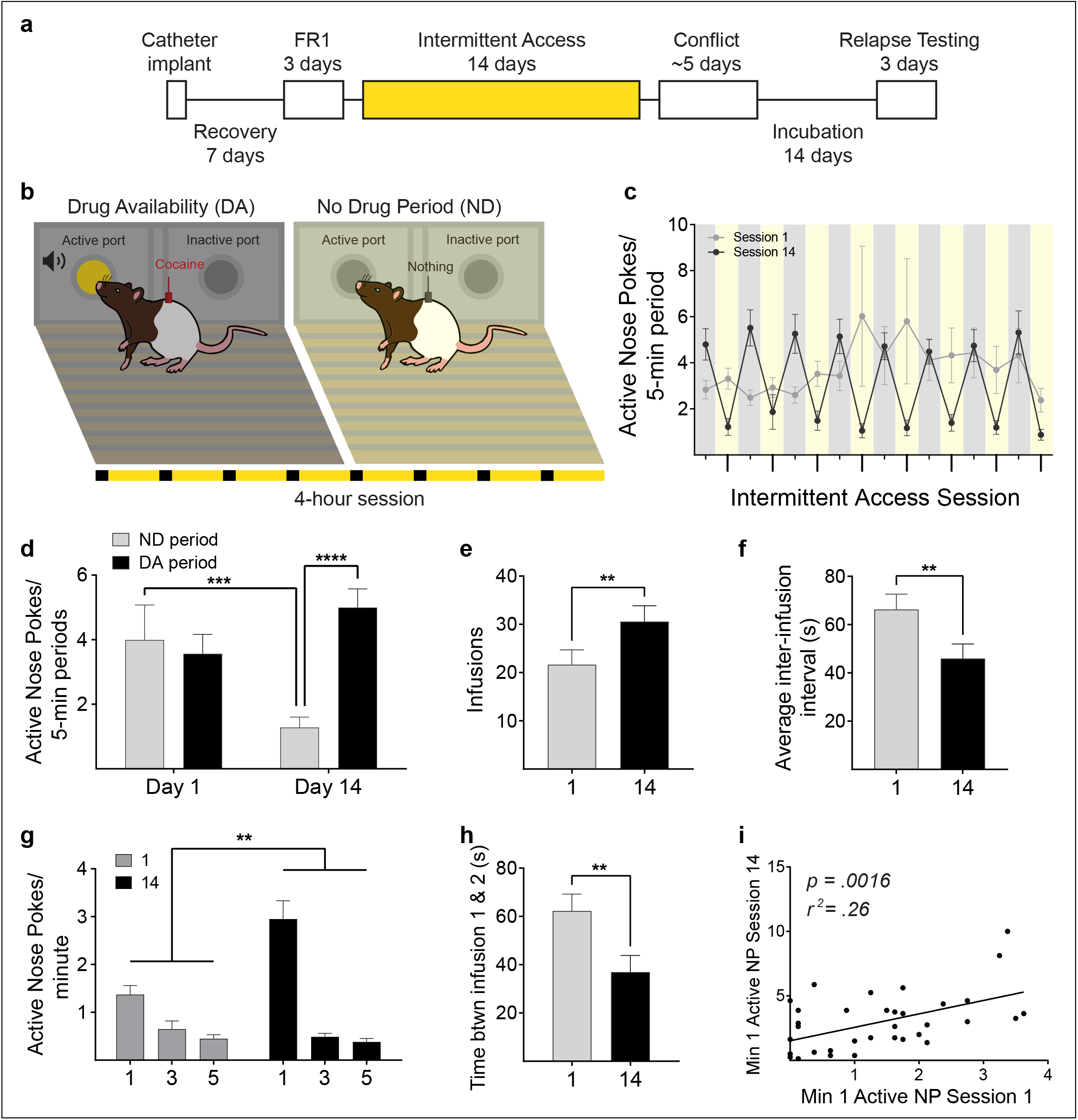
Intermittent access to cocaine promotes the development of rapid, binge-like intake that is under the control of drug availability state information. Intermittent access to cocaine self-administration paradigm and data. **a** Flow diagram of the experimental phases. Highlighted portion refers to the phase represented in the figure. **b** Schematic diagram of intermittent access showing the drug available period (left), no drug period (right), and pattern (bottom). **c** Rats learned to discriminate between the drug-available and no-drug periods across the 14 days. Gray and yellow boxes represent drug-available and no-drug periods, respectively. Gray and black circles represent session 1 and session 14, respectively. **d** Active nose pokes/5-min of drug available and no drug periods on sessions 1 and 14. Nose pokes decreased in the no-drug period across sessions. In session 14 nose pokes during the drug-available period were elevated compared to the no drug period. **e** Cocaine infusions increased from session 1 to 14. **f** Average inter-infusion intervals decreased from session 1 to session 14. **g** Within the drug-availability periods, active-nose pokes differed across the 5-min drug-availability period and this changed from session 1 to session 14. There was a main effect of minute, main effect of session, and a minute x session interaction. Specifically, active nose pokes within the 1st minute of the drug-available period were significantly elevated compared to the 3rd and 5th minute in session 1 and 14. As training progressed, rats made more active nose pokes during the 1st minute of drug availability periods. **h** The time between the 1st and 2nd cocaine infusions decreased from session 1 to session 14, demonstrating more vigorous binge-like drug taking with self-administration experience. **i** Initial binge-like drug use, defined as active nose pokes within the first minute of the drug available period, is positively correlated with binge-like drug use after 14-days of drug use. (n=35). Bars represent mean ± SEM. **p<0.01, ***p<0.001, ****p<0.0001.F(1,38)=1.104, p=0.3000).

Rats learned to discriminate between no-drug and drug-available periods across the 14 days (Figure 1D; no main effect of session F(1,34) = 0.9697, p=0.3317; main effect of availability period F(1.34) = 39.39, p<0.0001; session x availability period interaction F(1,34) = 19.72, p<0.0001). Active nose pokes during availability periods were significantly elevated above no-drug periods on session 14 (p<0.0001) but not session 1 (p=0.7676). Active nose pokes during the no-drug periods significantly decreased from session 1 compared to 14 (p=0.0005). Rats in the unpaired group failed to discriminate between no-drug and drug-available periods across the 14 days (Supplemental Figure 2C no main effect of session F(1,11) = 2.537, p=0.1395; no main effect of availability period F(1,11) = 2.269, p=0.1602; no session x availability period interaction F(1,11) = 2.220, p=0.1643). Paired rats significantly escalated their cocaine-intake across the 14 days (Figure 1E; t(34) =3.362, two-tailed, p=0.0019) and inter-infusion interval significantly decreased from session 1 to session 14 (Figure 1F; t(33)=3.079, two-tailed, p=0.0042) demonstrating rats took more cocaine more vigorously after 14-days of intermittent access. Given that our general sample included males and females, we next analyzed the data by sex. We found no sex differences in intermittent access behavior. Males and females similarly discriminated the drug availability and no drug periods (Supplemental Figure 3C; no main effect of sex F(1,35)=0.9735, p=0.3306) and self administered similar amounts of cocaine (Supplemental Figure 3D; no main effect of sex F(1,33)=0.002423, p=0.9610).

Within the drug-availability periods, active-nose pokes differed within the 5-min drug-availability period and this changed from session 1 to session 14 (Figure 1G; 2-way RM ANOVA, main effect of minute F(2,68) = 62.53, p<0.0001; main effect of session F(1.34) = 15.08, p=0.0005; minute x session interaction F(2,68)=18.59, p<0.0001). Specifically, active nose pokes within the first minute of the drug-available period were significantly elevated compared to the third and fifth minute in session 1 (1 v 3 p=0.0260; 1 v 5 p=0.0017) and session 14 (1 v 3 p<0.0001; 1 v 5 p<0.0001). Rats escalated their binge-like drug-taking, as active nose pokes during the first minute escalated from session 1 to session 14 (p<0.0001). Additionally, the time between the first and second cocaine infusions decreased from session 1 to session 14, demonstrating more vigorous binge-like drug taking with self-administration experience (Figure 1H; paired t-test, t(33) =3.506, two-tailed, p=0.0013). Intermittent access produced brief robust drug-taking that primarily occurred within the first min of the drug-available period, suggesting that the onset of the drug-availability state (box and house lights off) spurs rapid, binge-like cocaine use that escalated with drug-use experience. The pattern of responding during the drug availabile period was similar in males and females (Supplemental Figure 3F; no main effect of sex Active nose-pokes within the first minute of the drug-available period in session 1 correlated with active nose pokes within the first minute in session 14 (Figure 1I; F(1,33) = 11.84, p=0.0016; r2 = 0.2640). This suggests that initial binge-like drug use is associated with binge-like drug use after 14-days of drug use. This appears to be specific to behavior during intermittent access, as active nose pokes during the third day of acquisition (wherein a rat could take cocaine freely throughout the entire 4 hour session on a FR1 schedule) did not relate to active nose pokes during the drug-available period in either session 1 of intermittent access (Supplemental Figure 1D; F(1,33) = 0.6408, p=0.4292; r2 = 0.01905) or session 14 (Supplemental Figure 1E; F(1,33) = 0.9946, p=0.3259; r2 = 0.02926).

### Drug-availability state transitions spur rapid movement

Crossovers, a proxy for general locomotion, varied as a function of epoch (pre, during, and post drug-available period), session, and group (Figure 2B; 3-way interaction epoch x group x session F(2,44)=7.055, p=0.0022), increasing significantly during the DA period in paired (p=0.0010), but not unpaired (p=0.5745) rats. Making use of pose tracking analyses based on DeepLabCut (Figure 2C), we found that the transition to the drug-availability period almost instantly spurred psychomotor invigoration in paired rats. Distance traveled separated in 10-s bins (200s before to 50-s after drug-availability cue) varied as a function of group (Figure 2D; 2-way interaction bin x group F(7,154)=3.463, p=0.0018) and session (2-way interaction bin x session F(7,154)=2.622, p=0.0138). In the 20-30 sec after DA onset, Paired rats showed elevated distance traveled that significantly increased relative to unpaired rats across training days (Figure 2D,E; unpaired t test t(22)=1.74, p=0.048). Movement speed separated in 10-s bins also varied as a function of session and group (Figure 2F; 2-way interaction bin x group F(7,154)=2.356, p=0.0259; 2-way interaction bin x session F(7,154)=2.139, p=0.0427). Average speed in a group (paired or unpaired) varied as a function of session (2-way interaction group x session F(1,22)=5.507, p=0.0283). Speed increased rapidly in the ~30 sec after DA onset in the paired rats. Paired rats showed greater speed after DA onset that significantly increased relative to unpaired rats across training days (Figure 2F,G; unpaired t test t(22)=2.22, p=0.0185).

**Figure 2.**
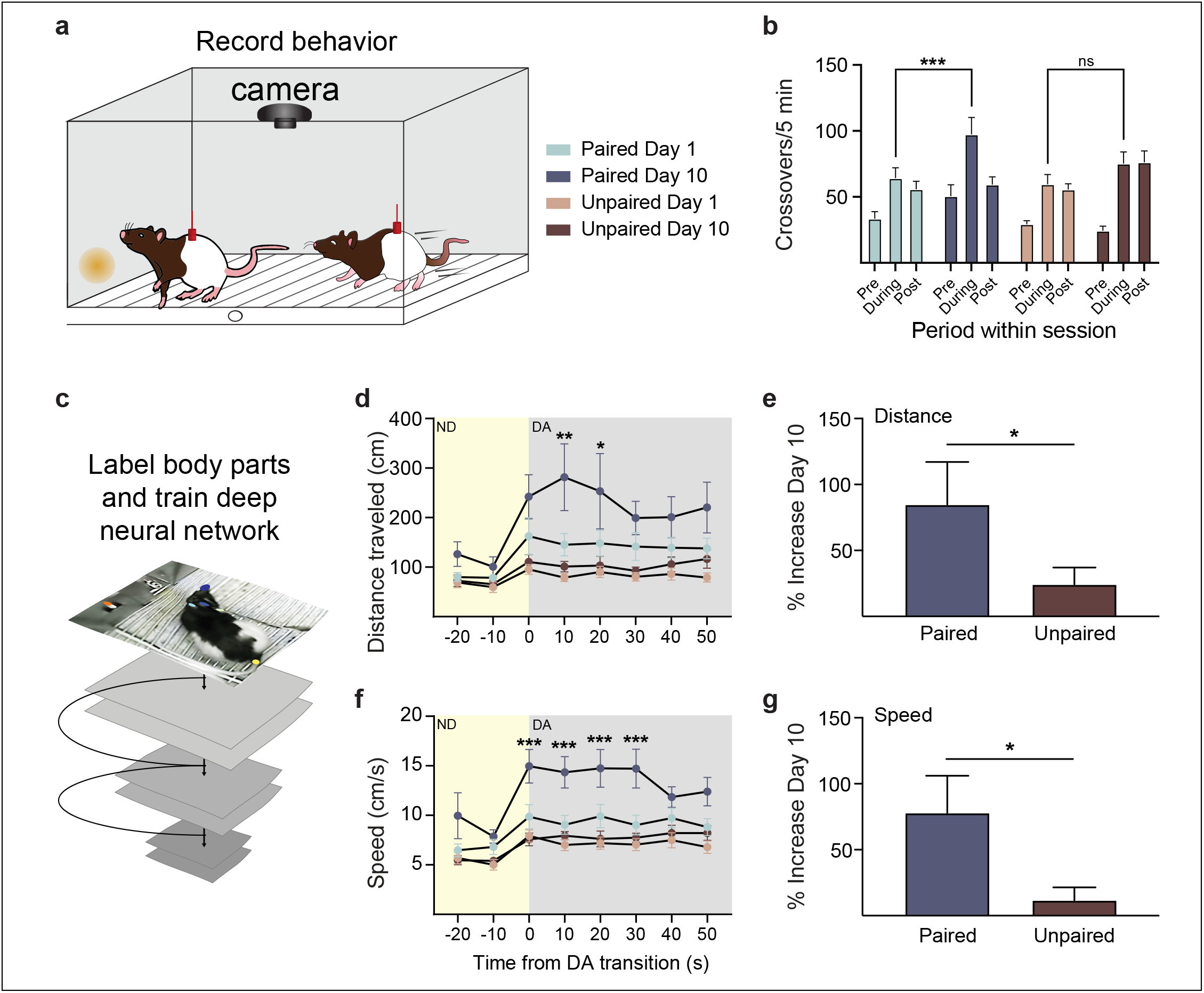
Drug availability states promote psychomotor sensitization when a drug seeking action-outcome contingency is intact. **a** Behavior is video recorded from cameras positioned above behavioral chambers, for hand scoring by experimenters and detailed analysis with DeepLabCut. **b** Intermittent access to cocaine led to increased chamber crossovers during the DA period in the paired (n=12) but not unpaired (n=12) rats. **c** Movement kinematics are interpolated from DLC-based pose estimation. **d-g** The transition to the drug-availability (DA) state spurs rapid psychomotor invigoration, with increased distance traveled **d-e** and speed **f-g**, in the first minute that sensitizes in paired compared to unpaired rats. Bars represent mean ± SEM. *p<0.05, **p<0.01, ***p<0.001.

### Binge-like drug use predicts persistent drug taking in the face of escalating cost

After intermittent access training, we used a conflict paradigm to assess drug taking persistence in the face of escalating cost. The grid floor in front of the nose-ports was electrified at increasing levels between sessions, such that rats had to overcome the cost (shock) to take cocaine (Figure 3B). For each individual rat, the cost was too high at a certain shock level and they remained in the shock-free zone, suppressing their drug taking behavior. Conflict ended with the shock level that resulted in cocaine intake at least two-thirds lower than behavior on the baseline session at 0 mA shock. Figure 3C displays the conflict effect of reduced drug seeking as shock increases. Across our subject pool, we found a distribution in shock levels rats were willing to persist through (Figure 3D). Males and females suppressed at similar shock levels (Supplemental Figure 3G unpaired t-test; t(33)=0.9490, two-tailed, p=0.3495). We found no relationship between cocaine intake at the end of continuous access pretraining and the levels of shock that rats later endure during conflict (Figure 3E; F(1,33)=0.9358, p=0.3404, r2=0.0276. We did find two predictors of the level of shock cost that rats were willing to persist through, which were behavioral measures taken after a history of intermittent cocaine binging: 1) that the amount of cocaine initially taken within the baseline shock level 0 session (Figure 3F; F(1,33) =6.019, p=0.0196, r2=0.1543), and 2) the amount of cocaine used in the last session of intermittent access (Figure 3G; F(1,33)=9.462, p=0.0042, r2=0.2228). This suggests that intermittent cocaine intake promotes cost insensitivity and higher levels of binge-like drug taking is associated with an increased motivation to persist through high costs to continue to take cocaine.

**Figure 3.**
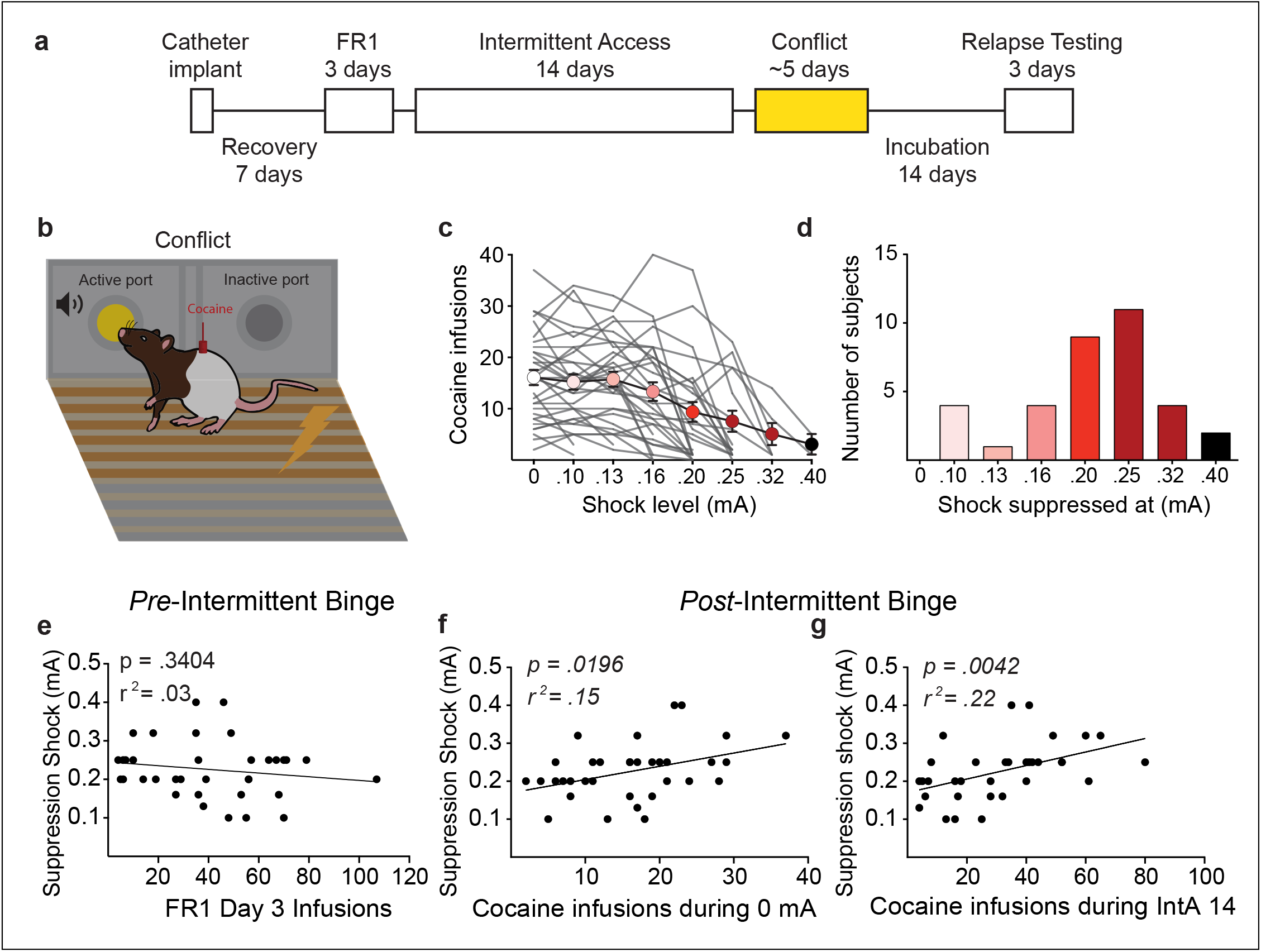
Binge-like cocaine self administration history predicts motivation to seek cocaine in the face of escalating cost. **a** Flow diagram of the experimental phases in which the phase represented in the figure is highlighted. **b** Schematic diagram of the conflict paradigm. **c** Average number of cocaine infusions decreases over escalating shock values. Bars represent mean ± SEM. **d** There was a distribution in shock levels rats were willing to persist through. **e** The amount of cocaine taken during the continuous access training phase, pre-intermittent binging, had no relationship with future shock cost the rats were willing to persist through. **f** The amount of cocaine initially taken within the baseline shock level 0 session predicted how much shock cost the rats were willing to persist through. **g** The amount of cocaine used in the last session of intermittent access was positively correlated with the shock intensity cost rats were willing to endure to continue taking cocaine. n=35.

### Drug cues interact hierarchically to spur drug seeking

After a 14-day incubation period, we assessed the influence of drug-associated cues to spur drug seeking in the face of cost, under extinction conditions. Within the single-cue test, the drug-delivery cue was non-contingently presented for 20s on a fixed 3-min interval during a 1-hr long session where ambient lights were off the entire time (Figure 4B). To make drug seeking responses, rats had to overcome a continuously-applied shock barrier in front of the nose ports. For relapse tests, the level of shock was set at 50% of what that individual rat suppressed to within the conflict phase, as in previous studies (Saunders et al. 2013). In the single-cue test, there were no differences between active nose pokes during the non-contingent drug-delivery cue presentation, active nose-pokes during the non-cued periods, and inactive nose-pokes (Figure 4C; F(2,48)=1.007, p=0.3727) suggesting that the drug-delivery cue on its own was not sufficient to trigger drug seeking. In the single-cue test for the unpaired group, there was also no difference in nose-pokes (Supplemental Fig. 2E; F(2,22)=2.053, p=0.1522).

**Figure 4.**
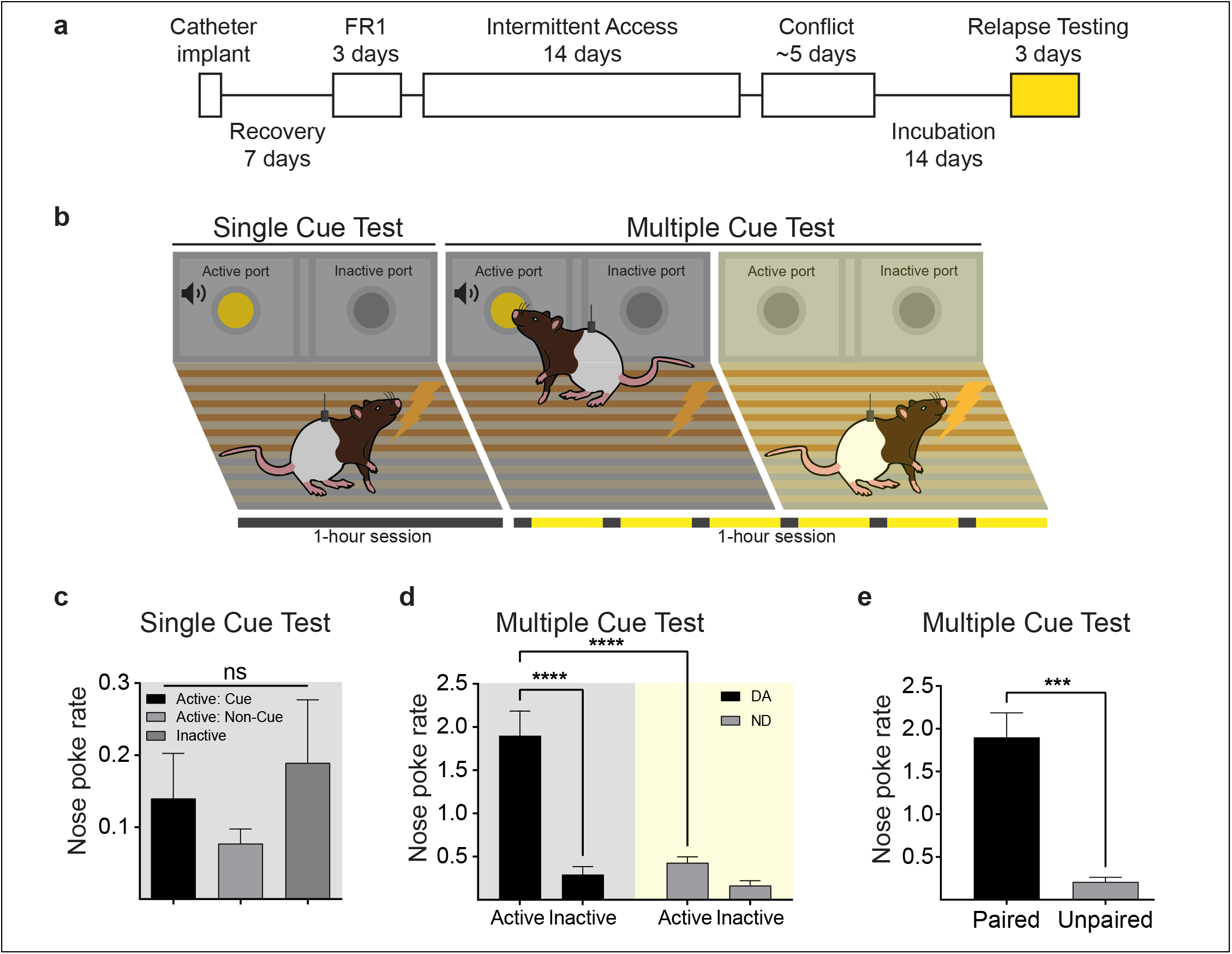
Drug-availability state transitions spur relapse in the face of cost and modulate the motivational value of proximal drug cues. **a** After a 14-day incubation period, we assessed the influence of drug-associated cues to spur drug seeking in the face of cost. **b** Schematic of the single cue test and multiple cue test. The single cue test consisted of a 20-s drug delivery cue that was presented non-contingently with a fixed 3-min ITI throughout a 1-hr session with the house lights off for the duration of the session. In the multiple cue test, the drug-availability cue (house lights going from on to off) was presented for 2-min every 8-min for a 1-hr long session. Within a drug-availability period, the 20-sec drug-delivery cue was non-contingently presented at 40s and 100s. **c** In the single cue test, active nose pokes during the non-contingent drug-delivery cue presentation did not differ from active nose-pokes during the non-cued periods (nose pokes/20sec). **d** In the multiple-cue test, non-contingent cue presentation of the drug-delivery cue spurred an increase in active nose pokes compared to non-cued baseline periods (nose pokes/2 min). **e** Only rats with a history of cocaine infusions paired with their drug seeking responses showed relapse in the multiple cue test. n=25. *p<0.05, **p<0.01.

This single-cue session was conducted entirely in darkness, so the rats never experienced the transition to the drug availability state, from light to dark. We hypothesized that global drug-availability state transitions may be an important mitigator of the motivational impact of this more proximal drug-delivery cue to spur drug seeking. We tested this with a multiple-cue test, wherein the drug-availability state (house lights going from on to off) was presented for 2-min every 8-min for a 1-hr long session (Figure 4B). Within a drug-availability period, the 20-sec drug-delivery cue was non-contingently presented every 40s. In the multiple-cue test, non-contingent cue presentation of the drug-delivery cue spurred an increase in active nose pokes compared to non-cued baseline periods (Figure 4D; multiple comparison’s test, p<0.0001, n=25), suggesting that information about drug availability gated the motivational impact of the drug-delivery cue to spur drug seeking. Nose poke rate was much higher in the paired group than the unpaired group during the drug available period of the multiple cue test (Figure 4E; unpaired t-test t(35) =4.059, p=0.0003). We found no sex differences in behavior during the multiple cue test (Supplemental Figure 3J; no main effect of sex F(1,23)=2.562, p=0.1231). Finally, we tested the ability of the drug delivery cue to reinforce nose poking on its own, in a 1-hr session conducted with houselights off. In contrast to noncontingent cue presentations in the single cue test, contingent cues reinforced robust responding in paired rats, compared to the unpaired group (Supplemental Figure 2H; unpaired t-test, t(35) =3.833, p=0.0005), with no sex differences (Supplemental Figure 3H; F(1,23)=4.064, p=0.0556).

### Individual variability in the influence of different types of drug-related cues to spur drug seeking

In the multiple-cue test, we saw individual differences in how rats responded to the drug availability state transition versus the delivery cue. A subset showed an increase in drug seeking in response to the drug-availability transition, before the delivery cue was presented. Another subset did not immediately respond to the drug-availability cue but showed elevated drug seeking during subsequent drug-delivery cue presentations. We categorized this heterogeneity using post-hoc criteria (Figure 5B). The group sizes and sex breakdown can be seen in Supplemental Figure 3K; 3 rats did not fit the criterion for either group. We set a criterion of at least a 2x increase in responding on the active nose port during the drug-delivery cue presentation compared to the periods where the drug-availability state was presented alone to be categorized in the proximal cue group (n=10). Rats in the proximal cue category increased their active nose pokes during the drug delivery cue presentations compared to baseline, but not during drug-availability periods in the absence of the delivery cue (Figure 5C; baseline vs. drug delivery cue p<0.0001, baseline vs. drug available p=0.5603). Figure 5D shows the pattern of responding on the active nose port for the proximal rats, (F(4,36)=7.577, p=0.0002) wherein there was an increase in responding to the drug-delivery cue presentations compared to baseline (p=0.0007 DD1, p=0.0033 for DD2), but did not alter their drug seeking in response to the drug availability state transition itself (p>0.05). This indicates that the impact of the drug-delivery cue to spur drug seeking is gated by the presence of drug availability state information, as these rats did not respond above non-cued baseline periods within the single cue test.

**Figure 5.**
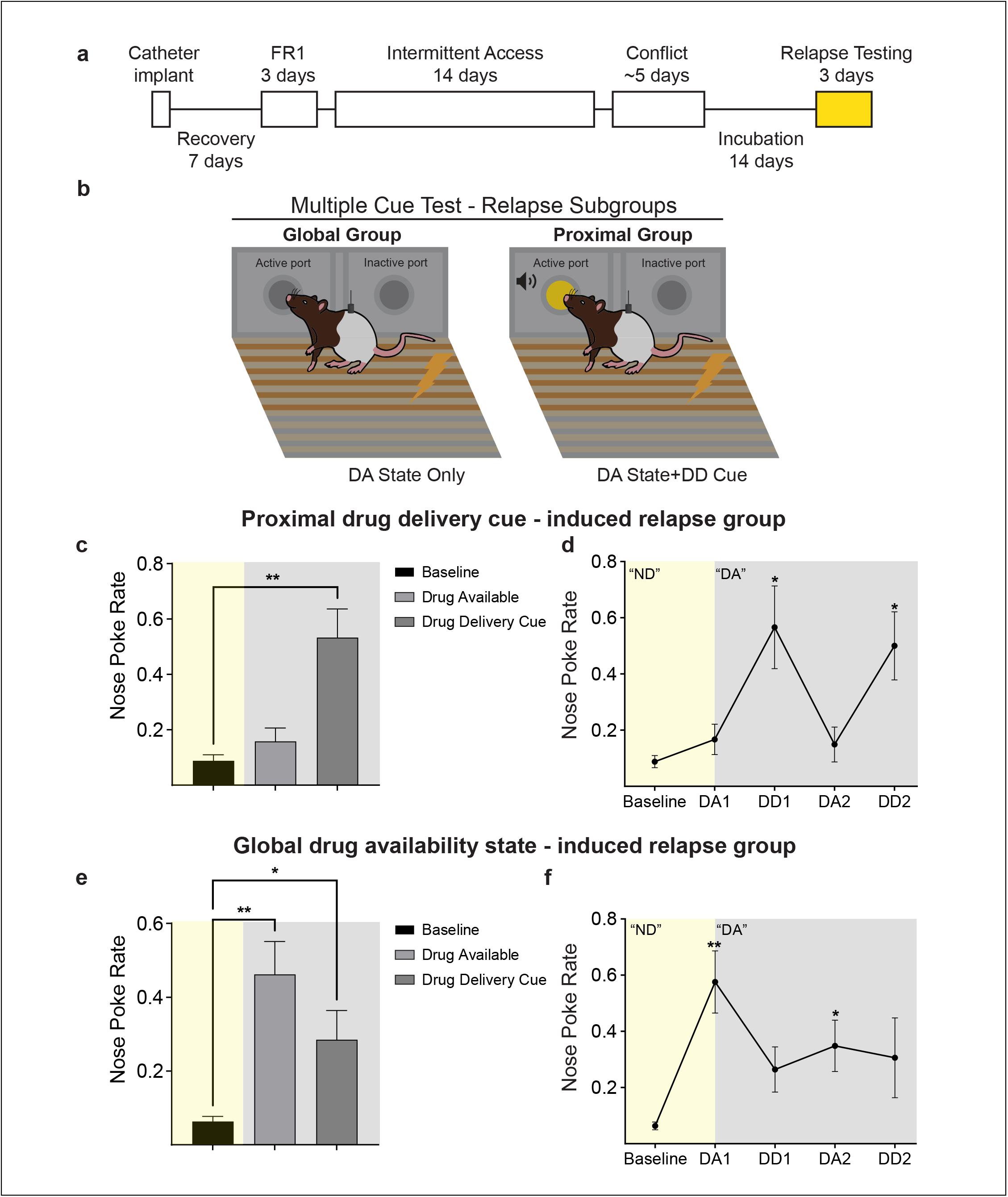
Individual differences in relapse triggers. **a** Flow diagram of the experimental phases in which the phase represented in the figure is highlighted. **b** Within the multiple cue test, relapse was preferentially triggered by proximal drug delivery cues or global drug availability state information in different subsets of rats. **c** Rats in the proximal group (n=10) increased their active nose poke rate during the drug delivery cue presentations, but not during the preceding drug-availability period (nose pokes/20sec). **d** The pattern of responding in the proximal group where nose pokes increase to the drug-delivery cues but not the drug-availability cue by itself. **e** For rats in the global group (n=12), nose poke rate increased immediately following the transition to the drug availability state. **f** The pattern of responding in the global group where nose pokes increase to the initial drug-availability cue. *p<0.05, **p<0.01.

To categorize rats that were driven to seek cocaine directly by the drug availability state transition, the global cue group (n=12), we set a criterion of at least 2x responding to the drug availability state onset (first 40 sec) compared to a no-cue baseline period. Rats in this group increased their active nose pokes compared to the baseline period during both the drug availability cue and the drug delivery cue periods (baseline vs. drug available p=0.0003, baseline vs. drug delivery cue p=0.0316). The global rats pattern of responding is observed in Figure 5F, (F(4,44) = 4.317, p=0.0049) wherein rats increased their active nose pokes at drug availability cue onset (p=0.0007). In an attempt to define a relapse subgroup predictor, we compared several variables in proximal and global rats. There were no differences between the subgroups in the number of cocaine infusions during intermittent access training, inter-infusion intervals during intermittent access training, level of shock reached during conflict, or conditioned reinforcement (Supplemental Figure 4; unpaired t tests; ps>0.05). Together this suggests that individual differences in sensory-guided relapse reflect an isolated phenotype from individual differences in binge-like cocaine intake and cost sensitivity.

## Discussion

Here, we modeled three key features of addiction: binge-like drug use, persistent drug seeking in the face of escalating costs, and the interaction of multi-cue relationships to spur relapse. We demonstrate that intermittent access produced robust drug taking invigorated by the transition to the drug availability state, which escalated across 14 days of self-administration. By the end of training, the bulk of infusions were taken in a rapid, binge-like fashion within the first minute of drug availability, similar to previous studies (Allain et al. 2018). We made use of a combination of experimenter and machine-learning guided analyses to quantify the movement profiles of self administering rats. This showed that the transition to drug availability states rapidly (i.e., within seconds) induces psychomotor sensitization, before cocaine infusions are taken. Critically, this state-induced movement invigoration was stronger compared to rats receiving rapid but passively delivered cocaine infusions. Thus, volitional drug intake reflecting intact action-outcome contingencies in the presence of dynamic state shifts is necessary for intermittent drug experience to promote exaggerated motivation. This result builds on the notion that intermittent access to drugs promotes sensitization to a greater degree than continuous long access (Allain et al. 2017; Algallal et al. 2020), and that drug-induced sensitization is under contextual control (Crombag et al. 2000).

### Hierarchically organized environmental control of drug seeking

Our results underscore the powerful role of the environment in regulating drug-related behaviors, emphasizing the influence of dynamic drug availability states on drug seeking. Drug availability spurred rapid movement invigoration and cocaine infusions during training. Critically, the presence of state transitions - the shift from an environment signaling drug unavailability to one signaling drug availability - was necessary for proximal drug-associated cues to induce relapse. This indicates that different forms of sensory information can interact in a hierarchical fashion to control drug seeking. We also found important differences between non contingent and contingent drug cue presentations in the promotion versus maintenance of drug seeking. Noncontingent cue presentations alone produced relatively weak relapse, compared to the ability of a drug cue presented after drug seeking responses to support conditioned reinforcement. Our results suggest that one potential reason for this is that the motivational value of non-contingently presented drug cues is strongly tied to the current state of the animal. Transitions to a state of drug expectation may “prime” individuals to be more vulnerable to the triggering influence of proximal drug cues. This distinction is notable because the majority of addiction studies in rodents examine relapse exclusively based on cue-reinforced responding, rather than non contingent cue presentations (Saunders et al. 2013; Venniro et al. 2020; Poisson et al. 2021).

Our relapse results could be partly explained by something akin to a Pavlovian to instrumental transfer effect, where the presentation of a Pavlovian conditioned stimulus invigorates instrumental reward-seeking actions (LeBlanc et al. 2012; Wassum et al. 2013). Given the modulating influence the drug availability state has on the motivational focal drug cues, these results also share elements of hierarchical or configural associative learning that is demonstrated in studies of contextual renewal and occasion setting (Chaudhri et al. 2008; Crombag et al. 2008; Fraser and Holland 2019; Valyear and Chaudhri 2020; Valyear et al. 2020). Our work in particular builds on previous studies demonstrating that drug-related contexts can regulate the rate of drug intake and the ability of discrete drug cues to reinforce drug seeking (Crombag and Shaham 2002; Sciascia et al. 2015; Valyear and Chaudhri 2020). However, with the transient nature of the drug availability period in our studies, we do not conceive of it as a classic contextual stimulus. Instead the drug availability period is a state, one that is toggled on and off during self administration sessions. Further work is needed to disambiguate these concepts to better understand state-dependent control of drug seeking and the intersecting influence of global and proximal drug cues.

### Binge-like cocaine intake promotes cost insensitivity in future drug seeking

We investigated the relationship between binge-like intake during intermittent access and the propensity to seek cocaine in the face of escalating costs, using a conflict procedure (Cooper et al. 2007; Barnea-Ygael et al. 2012; Saunders et al. 2013). We found that self administration training was predictive of seeking during conflict - rats taking more cocaine infusions at the end of intermittent access later endured higher shock levels to take cocaine. This suggests that rapid, intermittent cocaine intake facilitates cost-insensitive drug seeking, which may be a feature of compulsivity in addiction-like behavior. Continued drug seeking despite its unavailability has been conceptualized as an important aspect of addiction, and another proxy for compulsive or habitual behavior in animal models (Lüscher et al. 2020; Venniro et al. 2020; Poisson et al. 2021). In our studies, we did not see an increase in drug seeking responses during the no drug period, unlike some long access self administration studies (Deroche-Gamonet et al. 2004; Belin et al. 2009). Taken together, these results suggest that different behavioral correlates of compulsivity are not interchangeable and may instead reflect unique components of addiction.

### Individual differences in sensory-guided relapse trajectories

We find that state-level and proximal cocaine-related sensory information interacts hierarchically to guide drug seeking. Notably, however, considerable individual differences existed in which specific sensory elements promoted relapse across our subject pool. Some rats renewed drug seeking preferentially in response to state transitions - right after the drug availability period began. Other rats renewed seeking preferentially in response to the proximal drug delivery cue, following the transition to drug availability state. Thus, among individuals (rats in this case), there is variability in what type of environmental information is predominantly used to guide drug seeking behaviors.

Past work, including some of our own, demonstrated individual differences in the extent to which rats engage in cue-directed approach behavior (“sign tracking”), versus cue-evoked goal approach (“goal tracking”) predict drug cue-induced relapse and other addiction-like behaviors (Uslaner et al. 2006; Flagel et al. 2009; Saunders and Robinson 2010, 2013; Yager and Robinson 2013; Saunders et al. 2013). The potential relationship between sign/goal tracking variability and the individual differences we see here remains unknown. However, goal tracker rats are known to preferentially relapse in response to cocaine-associated contexts (Saunders et al. 2014), which could be akin to the drug-state induced relapse subgroup we found here. We found that individual rats’ binge-like self administration was correlated across intermittent access training, suggesting that an individual’s intake patterns set point was somewhat locked in, based on innate differences across subjects, rather than emergent from self administration experience. Notably, neither variability in cocaine intake rates during training nor cocaine seeking during conflict predicted which relapse subgroup a rat later fell into. This suggests that different facets of addiction-like behavior in our dataset - cost insensitivity and cued relapse vulnerability - represent somewhat independent traits. It remains unclear how to conceptualize this individual variability in relapse, but it may reflect broad inter-individual heterogeneity in decision-making strategies that intersect with the environment to produce different relapse phenotypes.

### Sex-based effects

Our studies included intact male and female Long Evans rats. We found no consistent evidence of sex differences. This is somewhat unexpected based on recent evidence that female rats show elevated motivation for cocaine during intermittent access self administration, resulting in more cocaine infusions, compared to males (Kawa and Robinson 2019; Algallal et al. 2020). The reasons for lack of sex differences here are unclear, but we note that the experimental protocols, including the length of pretraining acquisition and intermittent access training, differ substantially across the studies. Sex differences may be more apparent after longer periods of cocaine self administration. Previous studies also used other rat strains, including Sprague Dawleys. A recent report investigating variability in drug self administration across a broad sample of mouse strains showed that sex differences in drug seeking are not uniformly expressed, and instead depend heavily on the genetic background of animals (Bagley et al. 2022).

### Consideration of neural circuit mechanisms

Here we characterized multiple components of addiction-related behavior. Our results emphasize the moment-to-moment, dynamic nature of drug seeking motivation and complex interaction between the environment and individual cue-based behavioral variability. It will be essential to probe the brain mechanisms underlying our findings in future studies.

Dopamine plays a critical role in addiction (Robinson and Berridge 2001; Everitt and Robbins 2005; Collins and Saunders 2020). In SUD patients, an increase in striatal dopamine release to drug-paired cues is associated with an increase in drug craving and future relapse (Boileau et al. 2012). Blockade of dopamine receptors, conversely, attenuates drug-paired cue-induced craving (Franken et al. 2005). In preclinical studies, nucleus accumbens (NAc) dopamine signaling is critical for proximal drug-paired cues to promote and reinforce drug seeking action (Belin and Everitt 2008; Saunders et al. 2013). Furthermore, drug-paired cues evoke dopamine release (Ito et al. 2000; Aragona et al. 2009), and NAc dopamine signaling is required for drug-associated contextual information to renew cocaine seeking (Chaudhri et al. 2009; Saunders et al. 2014), suggesting that dopamine may track both proximal and state-level drug related information. Intermittent access to cocaine promotes sensitization of dopamine release and cocaine’s actions on the dopamine transporter in the NAc (Calipari et al. 2013, 2015; Kawa et al. 2019b). A key future direction will be to assess the impact of cocaine use on in vivo dopamine signaling dynamics during drug self administration and relapse wherein hierarchical cue relationships mitigate behavior. We would hypothesize that subsets of rats may show different dopamine signatures during drug seeking that could predict their unique relapse trajectories.

The basolateral portion of the amygdala (BLA) is implicated in the affective processing of environmental cues, including drug cues (Janak and Tye 2015; Wassum and Izquierdo 2015). This role is proposed to occur, at least in part, via downstream effects on NAc, where BLA neurons can modulate terminal dopamine release (Di Ciano and Everitt 2004; Jones et al. 2010). The BLA modulates the motivational impact of drug-paired cues by contextual information (Sciascia et al. 2015) and inhibition of the BLA-NAc circuit suppresses cue-related drug seeking (Stefanik et al. 2013; Puaud et al. 2021). The BLA is also implicated in state or model-based learning (Baxter and Murray 2002; Belova et al. 2008; Prévost et al. 2013; Saez et al. 2015; Puaud et al. 2021; Gostolupce et al. 2021). Further investigation is necessary to assess when, temporally, the BLA-NAc circuit contributes to drug seeking, and whether BLA neurons signal unique drug-related information, such as the value of availability states, that can be integrated downstream with NAc dopamine motivational signals. Other regions connected with the BLA, including the orbitofrontal cortex (OFC), have a key role in outcome- and state-based decision making (Wallis 2007; Schuck et al. 2016). In particular, OFC is important for discriminating state transitions, which is necessary for making appropriate decisions in dynamic environments (Bradfield et al. 2015; Stalnaker et al. 2021; Chan et al. 2021). Drug exposure may impair this process, resulting in persistent drug-seeking regardless of availability state or in the face of exaggerated costs (Lucantonio et al. 2012). The contribution of OFC circuits to our behavioral results is unknown but it will be important to explore how OFC signals respond to changing states in the context of intermittent drug self administration.

### Conclusion

Here, we describe studies that underscore the critical role of drug intake patterns and dynamic environments in regulating drug seeking motivation. Further, they suggest broad individual differences in sensory-guided relapse trajectories, which is an important consideration for the development of effectively targeted interventions for substance use disorders.

## Acknowledgements

This work was supported by National Institutes of Health grants T32-DA007234, F32-DA051138, and R00-DA042895. The authors declare no commercial or financial relationships that could be considered a potential conflict of interest.

## Author Contributions

VC and BS were responsible for experimental design. VC and KB conducted behavioral experiments. VC, KB, AW, and SS analyzed the data. VC, KB, and BS wrote the manuscript.

## Methods

### Subjects

All procedures were conducted in accordance with the National Research Council’s Guide for the Care and Use of Laboratory Animals and were approved by the University of Minnesota Institutional Animal Care and Use Committee. Intact male (~300-350g) and female (~250-300g) Long Evans rats (Envigo, Indianapolis, Indiana, USA) served as the subjects for these experiments (Intermittent Access and Conflict phases n=35, Cue-induced Relapse phase n=25; 12 additional rats were excluded due to catheter failure). Rats were initially group housed and then single housed post-catheter surgery and handled a week prior to the onset of the experiment. Experimentation took place during the light phase of their 12:12h light:dark cycle. Free access to water and food chow was provided in their home cage.

### Intravenous Catheter Surgery

Catheter back ports were purchased from Instech (Plymouth Meeting, PA, USA) and prepared with 0.51 mm inner/0.94 mm outer diameter silastic tubing at 11.5 cm length with a silicone ball at 2.5 cm length. Rats were anesthetized with ketamine hydrochloride (10mg/ kg; i.p.) and xylazine (10mg/kg; i.p.). Intra-jugular catheter placement was conducted as described previously (Saunders and Robinson 2010; Saunders et al. 2014). Briefly, after securing the end of the indwelling tubing within the right jugular vein, the connecting port was secured on the back and protected with a fitted cap (Instech). Following surgery, catheters were flushed daily with 0.1 ml of Gentamicin sulfate (5mg/ml; Vedco, MO) to prevent infection and line blockage. Catheter patency was tested prior to and at the end of each experimental phase (~2 weeks) with 0.1 ml methohexital sodium (10mg/ml, i.v.). Only patent rats, determined by ataxic response 3-5sec post-injection, were included in the analyses.

### Cocaine self-administration: Acquisition

One week after catheter surgery, rats learned to self-administer cocaine in three continuous-access sessions. Med Associates chambers were outfitted with two ports on the left side that served as the active and inactive nose-ports (counterbalanced between subjects). A nose poke into the active nose port resulted in an IV infusion of cocaine (0.4mg/kg/infusion in 50 ul sterile saline, delivered over 3.2 sec; Boynton Pharmacy, Minneapolis, MN) on a fixed-ratio (FR)1 schedule of reinforcement with a 20-sec timeout period, during which nose-pokes had no consequence. Active nose pokes also resulted in the illumination of the nose port and 75dB white noise for 20 sec (the drug delivery cue). Nose pokes in the inactive nose port had no consequences. These sessions were conducted with the ambient lighting in the chambers (house and box lights) turned off.

We included an unpaired control group to examine the effects of rapid, intermittent cocaine exposure that was not the result of a specific action-outcome contingency. This was achieved by unpairing the action (nose poke) from the outcome (cocaine infusion) and the Pavlovian relationship (drug delivery cue and cocaine infusion). We retained the relationship between the drug availability state and cocaine exposure. The rate, pattern and magnitude of cocaine infusions and cue deliveries was matched for sex and session to the average levels achieved by the paired self administration cohort. For pretraining in unpaired rats (n=12), a 20-sec cue consisting of illumination of a nose port light and white noise - akin to the drug delivery cue in paired rats - was presented non-contingently throughout a 4-hour session on a variable interval schedule based on sex (average female ITI (min) days 1-3: 6.0, 7.1, 6.9; average male ITI (min) days 1-3: 7.7, 6.5, 7.3). Rats also received non-contingent cocaine infusions (3.2-sec infusion, 0.4 mg/kg) via an intravenous catheter throughout the session with the same variable interval schedule based on sex. The number of infusions and cue presentations were matched to the average from the paired group of that day based on sex (female days 1-3: 40, 34, 35; male days 1-3: 31, 37, 33). In this way, the cue was not associated with cocaine delivery, and nose pokes did not earn cocaine.

### Cocaine self-administration: Intermittent Access

After acquisition, rats went through 14-days of intermittent access similar to previous studies (Zimmer et al. 2012; Calipari et al. 2013; Kawa et al. 2019b). Within each 4-hour session, there were 5-min periods wherein active nose pokes resulted in a cocaine infusion (0.4mg/kg/infusion in 50 ul in 3.2 sec) that coincided with drug delivery cue (3.2 sec) on a FR1 schedule of reinforcement. In this case, the timeout period lasted only 3.2 sec, coinciding with the duration of each infusion. These drug-available (DA) periods were signaled by the extinguishing of the house and box lights for the entire 5-min period. Following the drug-available period, the house and box lights illuminated, signaling the 25-min no-drug (ND) period, wherein active nose-pokes had no consequence. This pattern (5-min DA and 25-min ND periods) repeated 8 times for a 4-hr session. Inactive nose-pokes had no consequences. wwFor unpaired rats, within each session a 5-minute drug exposure period (signaled by ambient lights off) was followed by a 25-minute drug unavailability period (lights on) which patterned 8 times throughout the session, as with paired rats. In the drug exposure period, rats received non-contingent cocaine infusions (3.2 s, 0.4 mg/kg) on a variable interval schedule based on sex (average female ITI (min) days 1-14: 3.0, 2.6, 2.0, 2.2, 1.9, 1.9, 1.8, 1.7, 1.8, 1.9, 1.6, 1.5, 1.3, 1.5; average male ITI (min) days 1-14: 1.6, 1.5, 1.7, 1.4, 1.4, 1.2, 0.9, 1.4, 1.1, 1.4, 1.1, 1.3, 1.2, 1.1). Throughout the entire session, rats received non-contingent 3.2-sec cue presentations on a variable interval schedule based on sex (average female ITI (min): 17.7, 15.3, 12.0, 12.9, 11.3, 11.7, 10.8, 10.3, 11.1, 11.6, 9.6, 9.2, 8.0, 9.0; average male ITI (min): 9.5, 9.2, 10.1, 8.2, 8.4, 7.2, 5.5, 8.1, 6.7, 8.2, 6.7, 7.7, 7.2, 6.7). The number of infusions and cue presentations they received was the average from the paired group of that day based on sex (female days 1-14: 14, 16, 20, 19, 21, 21, 22, 23, 22, 21, 25, 26, 30, 27; male days 1-14: 25, 26, 24, 29, 29, 33, 44, 30, 36, 29, 36, 31, 33, 36). In this way, unpaired rats received cocaine infusions at the same average rate during the drug exposure period across sessions, compared to those volitionally taken by paired rats.

### Cocaine self-administration: Conflict

After 14 days of intermittent access rats underwent a conflict phase (Cooper et al. 2007; Saunders et al. 2013). For each 1-hr session, 2/3 of the grid floor in front of the nose ports within the chamber was continuously electrified at escalating levels of shock between sessions (0, 0.13, 0.16, 0.2, 0.25, 0.32, 0.4 mA). The back 1/3 of the grid floor was not electrified, allowing rats to avoid shock. Rats received increasing shock intensities on successive sessions until the number of cocaine infusions was reduced at least 2/3 below baseline levels determined by the 0mA shock-level session. The house and chamber lights were extinguished throughout the entire 1-hr session. An active nose-poke resulted in a cocaine infusion (0.4mg/kg/ infusion in 50 ul over 3.2s) that coincided with the drug delivery cue for 20 sec. There was a 20-sec timeout period after each cocaine infusion wherein additional active nose pokes had no further consequence. Inactive nose pokes had no consequences.

For unpaired rats, conflict was similar. Current increased over 6 1-hr sessions (0, 0.1, 0.13, 0.16, 0.2, 0.25 mA). The number of sessions was determined by the average amplitude at which rats in the paired group suppressed drug seeking behavior, as defined by 2/3 reduction in nose pokes from a no shock baseline. Rats received non-contingent cocaine infusions unpaired from non-contingent cue presentations at a variable interval schedule based on sex (average female ITI (min) days 1-6: 3.2, 3.3, 3.0, 4.0, 5.0, 6.7; average male ITI (min) days 1-6: 4.0, 4.0, 4.3, 5.5, 7.5, 7.5). Number of cocaine infusions and cue presentations was the average from the paired group of that day based on sex (female days 1-6: 19, 18, 20, 15, 12, 9; male days 1-6: 15, 15, 14, 11, 8, 8).

### Cue-induced drug seeking

Following 14 days of abstinence, rats underwent tests for cue-induced relapse, wherein we assessed the influence of non-contingent cue presentations on drug-seeking in the face of cost. During these sessions, 2/3 of the grid floor was electrified at the shock intensity 50% to the level of each rat’s individual suppression shock level. For example, if a subject suppressed in the conflict phase at 0.4 mA, 0.2mA was used for the cue-induced drug seeking tests. To assess drug-seeking motivation in absence of drug intoxication, no cocaine was delivered during these tests. In the single-cue test, the drug-delivery cue (illumination of the active nose-port and 75 dB white house) was non-contingently presented for 20 sec with a fixed 3-min intertrial interval (ITI) for 1-hr. The house and box lights were extinguished throughout the entire single-cue test. For the multiple-cue test, (order of single and multiple was counterbalanced between subjects), the drug-availability state (extinguished house and box lights) was presented for 2 min on a fixed 8-min ITI, in a 1-hr test session. Within each 2-min availability period, the 20-sec drug-delivery cue (illumination of the nose port and 75 dB white noise) was presented twice (at 40 seconds and 100 seconds). Nose pokes had no consequences. For relapse tests in unpaired rats, shock was set at 0.13 mA. Both cue tests were run twice, once with the drug delivery cue (nose port light and white noise) and once with a neutral cue (other nose port light and white noise), to examine spontaneous responding.

### Conditioned Reinforcement

To assess the reinforcing value of the drug-delivery cue, rats underwent a conditioned reinforcement test. During this 1-hour session, active nose pokes resulted in a 3.2-sec presentation of the drug-delivery cue but no cocaine infusion. Inactive nose poke did not result in any consequence. Box and house lights were extinguished during the entire session. Unpaired rats also underwent conditioned reinforcement, once with the drug delivery cue (nose port light and white noise) and once with a neutral cue (other nose port light and white noise).

### Video scoring

Med Associates chambers were fitted with overhead cameras (Vanxse CCTV 960H 1000TVL HD Mini Spy Security Camera 2.8-12mm Varifocal Lens Indoor Surveillance Camera) for a top-down video recording of self-administration behavior. We quantified locomotor activity during intermittent access via visual examination of these videos, similar to previous studies (Carr et al. 2020). The chamber was divided into four equal quadrants and locomotor activity was determined by counting the number of crossovers, defined as when the animal’s back port fully crossed a line dividing the quadrants. Behavior was quantified for 3 separate epochs: 5 min before (pre), 5 min during, and 5 min after (post) the drug availability period. This was done for sessions 1 and 10.

### DeepLabCut-based pose tracking

Markerless tracking of animal body parts was conducted using the open-source DeepLabCut (DLC) Toolbox (Mathis et al. 2018) and analysis of movement features based on these tracked coordinates was conducted in Matlab R2020b (Mathworks). All DLC analysis was conducted on a Dell G7-7590 laptop running Windows 10 with an Intel Core i7-9750H CPU, 2.60Ghz, 16 GB RAM, and an NVIDIA GeForce RTX 2080 Max-Q 8GB GPU. DeepLabCut 2.1.10 was installed in an Anaconda environment with Python 3.7.7 and Tensorflow 1.13.1. Videos (944 x 480 resolution) were recorded with a sampling frequency of 30 frames per second using a TIGERSECU Super HD 1080P 16-Channel DVR system.

### DeepLabCut Model

A previous DLC network trained for 500,000 iterations on 1700 manually labeled frames extracted from 17 videos (16 different animals, 9 different recording sessions) was updated for the current study. An additional 370 frames from this experiment were extracted (25 frames each uniformly extracted from 14 videos of 12 different animals), and 20 manually extracted frames for additional examples where the animal was rearing. Body parts were manually labeled in all frames for the nose, eyes, ears, center of head, catheter port, and tail base. Features of the environment were also labeled manually, including the 4 corners of the apparatus floor and the nose ports.

Labeled frames were split into a training set (95% of frames) and a test set (5% of frames) and trained using the training set for 500,000 iterations. An additional 267 outlier frames were extracted from 7 videos over 2 iterations. For each iteration, the network was retrained from the default Resnet50 model for 1,030,000 iterations using a newly generated 95% train–test split. The model was evaluated by comparing the labels acquired from the trained network on the training and test sets with the manual user labels. Evaluation gave an error of 2.74 and 3.52 pixels for the training and test sets. Using a p-cutoff of 0.85 error was reduced to 2.63 pixels for the training set and 3.08 pixels for the test set. This model was then used to analyze videos from 24 animals (12 paired and 12 unpaired) for two separate intermittent access sessions (session 1 and session 10). Data for 1 session was re-analyzed for 2 animals after a further iteration. Pixel errors for this iteration were 2.65 and 2.93 pixels for the training and test datasets, 2.55 for the training set and 2.88 for the test set with a p-cutoff of 0.85.

## Data Analysis

### Self-administration behavior

Nose pokes and cocaine infusions were the primary behavioral output measures. Data were processed with Microsoft Excel (Richmond, WA). Statistical analyses were conducted with GraphPad Prism (La Jolla, CA). For all hypothesis tests, the α level for significance was set to p<0.05. Data were analyzed with mixed effects ANOVA, Pearson correlation, and linear regression. Post-hoc comparisons and planned t-tests were used to clarify main effects and interactions.

### Video analysis

#### Hand-scored locomotion data

Chamber crossover data was analyzed in GraphPad Prism with an ANOVA or mixed effects analysis comparing epoch (5-min bins), group (unpaired or paired), and session (1 or 10).

#### DeepLabCut data

DLC data were analyzed with MATLAB and GraphPad Prism. All data for four animals, two from each group (paired/unpaired) were excluded from analysis due to the commutator obscuring the view of the active nose port in the recorded video. The x,y co-ordinates of fixed environment features were averaged across all frames where the likelihood (confidence-interval) value was > 0.7. The coordinates of the corners of the apparatus floor were used to calculate a pixel to cm conversion for each animal and session. This was calculated by taking the average pixel-to-cm ratio of all 4 sides and the two 2 diagonal measurements of the floor. For instances where not all points were visible in the video frame, only the available measurements were included in the average.

Frames where likelihood values for the body part x,y coordinates were < 0.7 were removed from the data before further processing. For all measures except movement speed, missing data for periods of <1s due to low confidence was replaced using a moving average between the bordering coordinates before subsequent analysis. Movement speed and distance traveled were calculated using the x,y coordinates of the Catheter port. Speed was calculated from frame to frame using the formula: distance (cm)/time (s). Movement speed for each epoch was smoothed with a moving mean average over a window of 0.1s, using the MATLAB function smoothdata. Distance traveled was calculated as the distance between x,y coordinates from frame to frame and converted to cm using that animal’s pixel-to-cm ratio. Data were analyzed for 3 separate epochs: 5 min before (pre), 5 min during, and 5 min after (post) the drug available period. We also analyzed a shorter time window from 20s before to 60s after the drug availability period onset with data binned in 10s increments. Analysis windows where >40% of coordinate data was missing for the catheter port were considered invalid and the measures dependent on these coordinates were not included in the analysis.

---

**Supplemental Figure 1.**
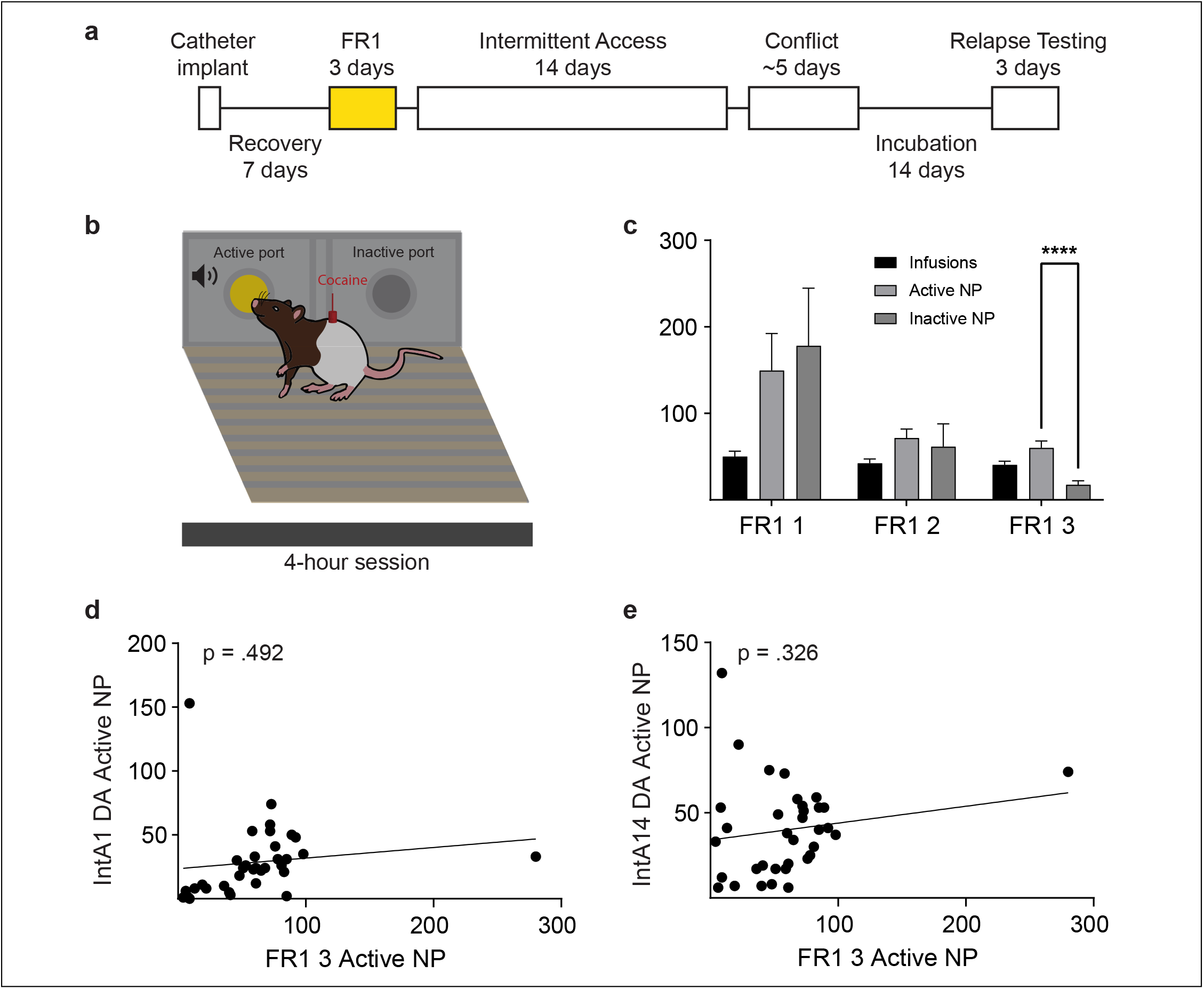
**a** FR1 pretraining data. **b** Schematic of FR1 pretraining where a nose poke in the active port resulted in a cocaine infusion paired with a drug-delivery cue. **c** Three days of FR1 pretraining, during which rats learned to discriminate between active and inactive nose pokes. Bars represent mean ±SEM. Active nose pokes on the final day of acquisition did not correlate with active nose pokes on **d** day 1 or **e** 14 of intermittent access. n=35. ****p<0.0001.

**Supplemental Figure 2.**
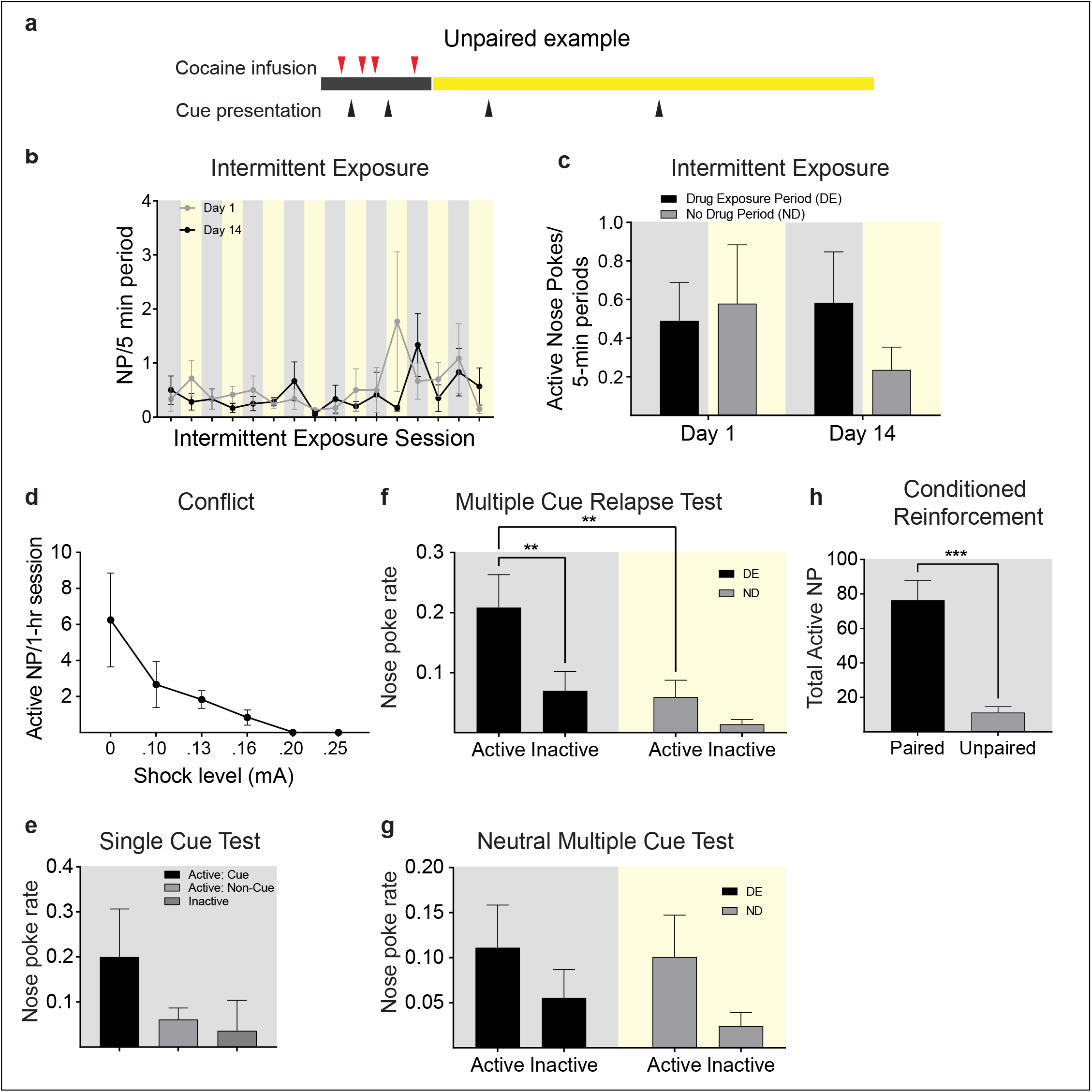
Unpaired cocaine exposure data. **a** A schematic example of the schedule for a drug-exposure period (lights off) and no-drug period (lights on). Cocaine infusions were delivered non-contingently in the drug-exposure period while drug-delivery cue presentations were delivered throughout both periods. **b** Rats in the unpaired group did not make nose pokes during the drug-exposure or no-drug periods across the 14 days. Gray and yellow boxes represent drug-exposure and no-drug periods, respectively. Gray and black circles represent session 1 and session 14, respectively. **c** Active nose pokes/ 5-min of drug exposure and no drug periods on sessions 1 and 14 of the unpaired group. Nose pokes did not change in regard to period or session. **d** Average number of nose pokes decreases over escalating shock values in the unpaired group. **e** In the unpaired single cue test, active nose pokes during the non-contingent drug-delivery cue presentation did not differ from active nose-pokes during the non-cued periods. **f** In the unpaired multiple-cue test, non-contingent cue presentation of the drug-delivery cue spurred a small increase in active nose pokes compared to non-cued baseline periods. **g** When the drug-delivery cue was changed to a neutral cue in the unpaired multiple-cue test, non contingent delivery of the neutral cue failed to elicit differences in behavior between periods. **h** Active nose pokes during conditioned reinforcement for a drug delivery cue presentation were significantly higher in the paired group than the unpaired group. Bars represent mean ± SEM. n=12.

**Supplemental Figure 3.**
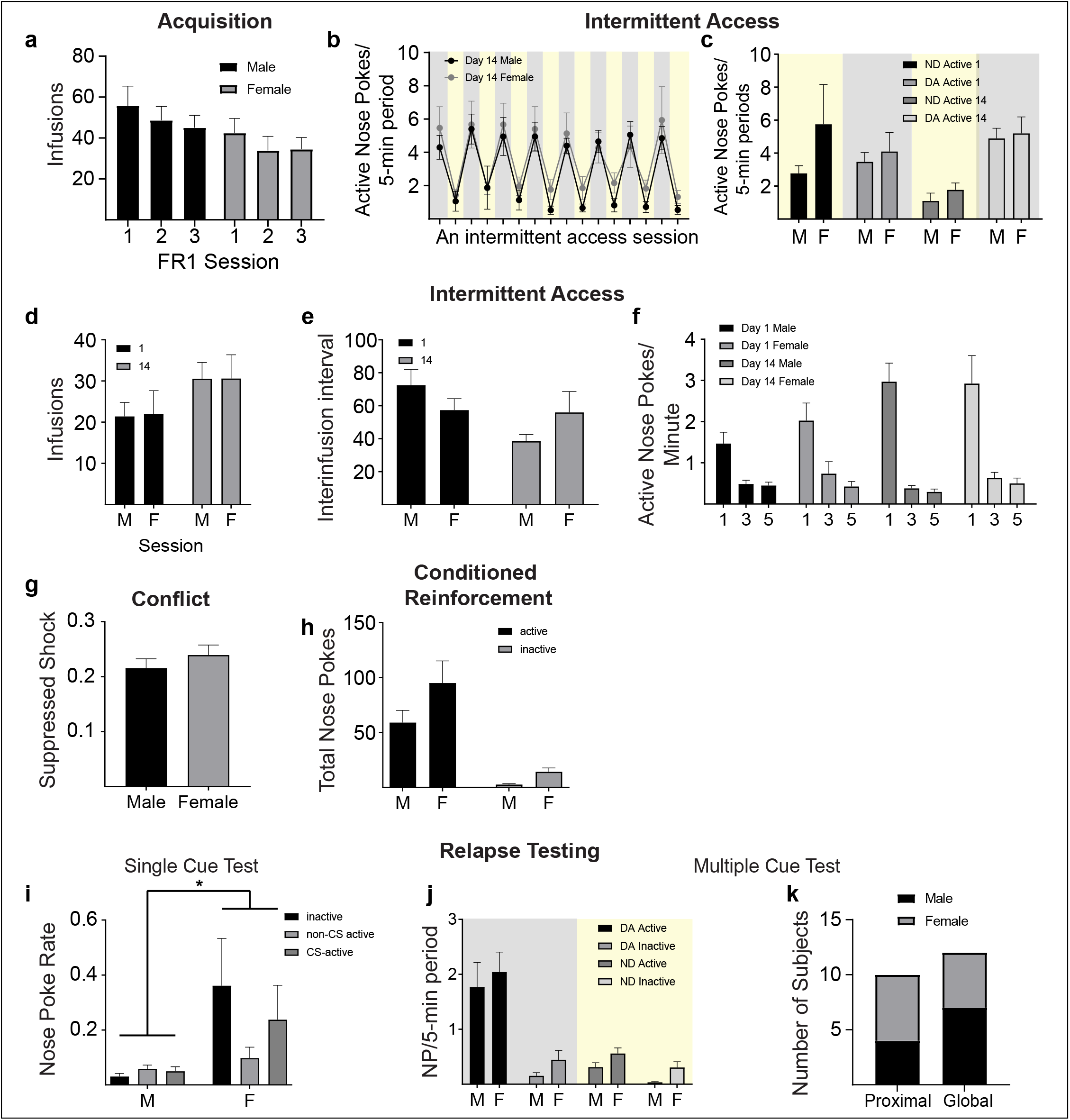
No clear sex differences emerged. **a** Males and females used similar amounts of cocaine throughout FR1 pretraining. **b,c** Male and female rats similarly learned to discriminate between the drug-available and no-drug periods across 14 days of intermittent access training. **d** Cocaine intake escalated across training similarly in males and females. **e** Males and females had similar inter-infusion intervals throughout intermittent access. **f** Within the drug-availability periods, active-nose pokes did not differ by sex across the 5-min drug-availability period on sessions 1 and 14. **g** Males and females suppressed cocaine seeking behavior at similar shock thresholds. **h** No sex differences emerged during conditioned reinforcement behavior. **i** There was a main effect of sex during the single cue test. **j** There were no sex differences in behavior during the multiple cue test. **k** The sex breakdown was similar within the two relapse subgroups. Proximal group: 4 males and 5 females. Global group: 7 males and 5 females. Bars represent mean ± SEM. *p<0.05.

**Supplemental Figure 4.**
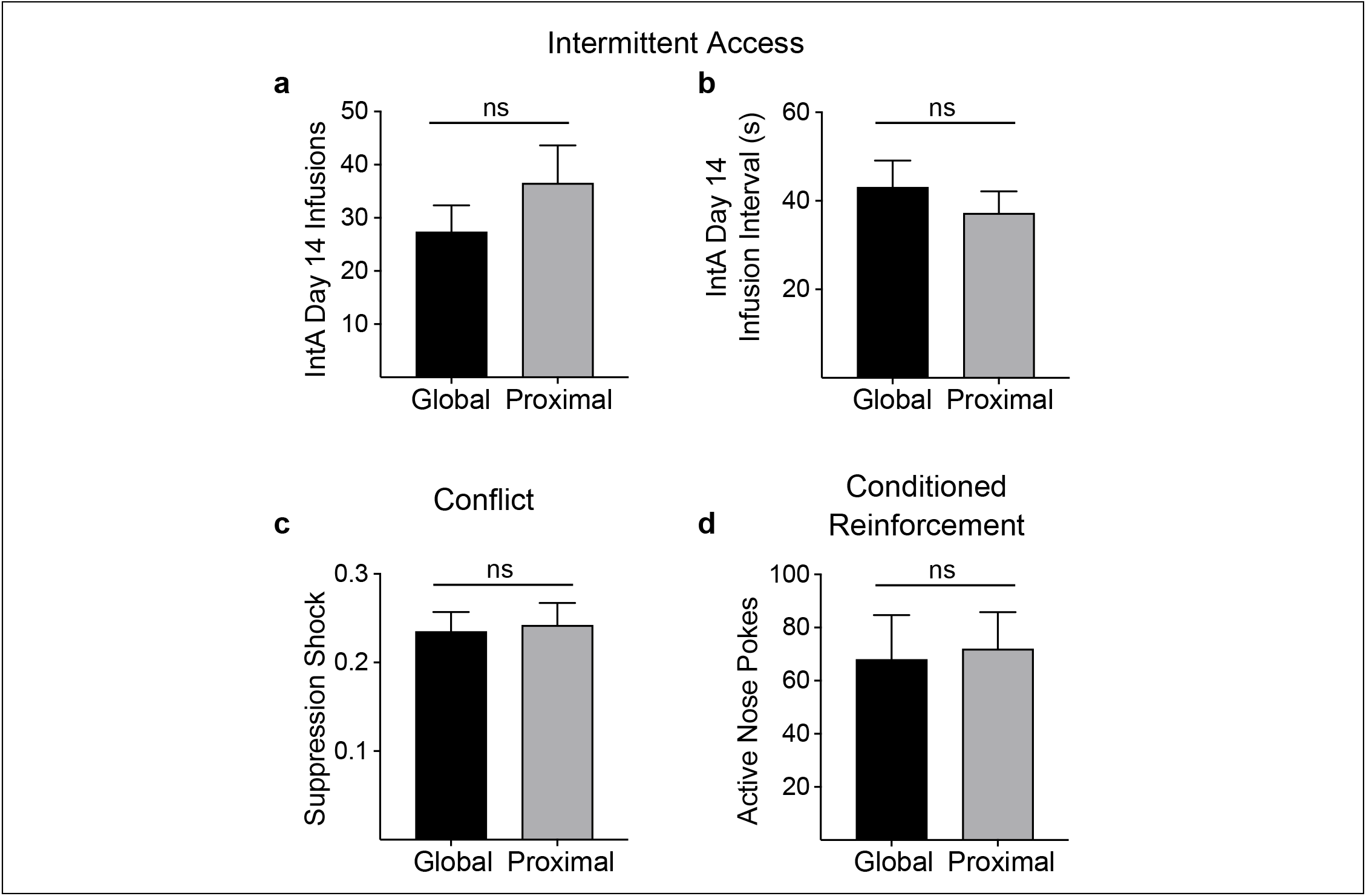
Self administration variables do not predict relapse subgroups. We found no differences in **a** the number of cocaine infusions during intermittent access training, **b** inter-infusion intervals during intermittent access training, **c** level of shock reached during conflict, or **d** conditioned reinforcement between the global (n=12) and proximal (n=10) relapse subgroups. Bars represent mean ± SEM.

